# Monoallelic antigen expression in trypanosomes requires a stage-specific transcription activator

**DOI:** 10.1101/2021.05.06.442931

**Authors:** Lara Lopez Escobar, Benjamin Hänisch, Clare Halliday, Samuel Dean, Jack Daniel Sunter, Richard John Wheeler, Keith Gull

**Author notes:** Correspondence and requests for materials should be addressed to Richard Wheeler & Jack Sunter &.

## Abstract

Monoallelic expression of a single gene family member underpins a molecular “arms race” between many pathogens and their host, through host monoallelic immunoglobulin and pathogen monoallelic antigen expression. In *Trypanosoma brucei*, a single, abundant, variant surface glycoprotein (VSG) covers the entire surface of the bloodstream parasite^1^ and monoallelic VSG transcription underpins their archetypal example of antigenic variation. It is vital for pathogenicity, only occurring in mammalian infectious forms^1^. Transcription of one VSG gene is achieved by RNA polymerase I (Pol I)^2^ in a singular nuclear structure: the expression site body (ESB)^3^. How monoallelic expression of the single VSG is achieved is incompletely understood and no specific ESB components are known. Here, using a protein localisation screen in bloodstream parasites, we discovered the first ESB-specific protein: ESB1. It is specific to VSG-expressing life cycle stages where it is necessary for VSG expression, and its overexpression activates inactive VSG promoters. This showed monoallelic VSG transcription requires a stage-specific activator. Furthermore, ESB1 is necessary for Pol I recruitment to the ESB, however transcript processing and inactive VSG gene exclusion ESB sub-domains do not require ESB1. This shows that the cellular solution for monoallelic transcription is a complex factory of functionally distinct and separably assembled sub-domains.

---

The single VSG gene is expressed from a specialised bloodstream form telomeric expression site (BES), along with 4 or more expression site associated genes (ESAGs)^4–6^. Switching VSG is achieved by switching to transcription of one of several different telomeric BESs^5^ or replacement, by recombination, of the VSG in the active BES with one of the ∼2500 VSG gene and pseudogene variants elsewhere in the genome^7^. Only the active BES is found in the Pol I-containing^2^ specialised, non-nucleolar, nuclear structure called the ESB. The ESB is only present in bloodstream form parasites^3^, despite procyclic forms (in tsetse fly) also employing Pol I-dependent transcription of their invariant surface coat (procyclin). Elegant biochemical candidate approaches and genetic screens of VSG expression have revealed the importance of epigenetic silencing^8^, telomere^9–12^ and chromatin factors^13–20^ and SUMOylation^21,22^. VEX proteins, required for exclusion of the inactive BESs^23,24^, associate the single active BES with the Spliced Leader (SL) array^25^ chromosomal locations. These contain the repetitive genes encoding a sequence which, after transcription and processing, is added to every trypanosome mRNA^26^. Hence, in addition to other properties, VEX proteins link an ESB located exclusion phenomenon to an active VSG gene abundant mRNA processing capability. Notwithstanding these advances, bloodstream-specific factors (Fig. 1A, Extended Data Table 1) remain elusive and the statement that “No ESB-specific factor has yet been identified”^27^ still holds true.

**Fig. 1.**
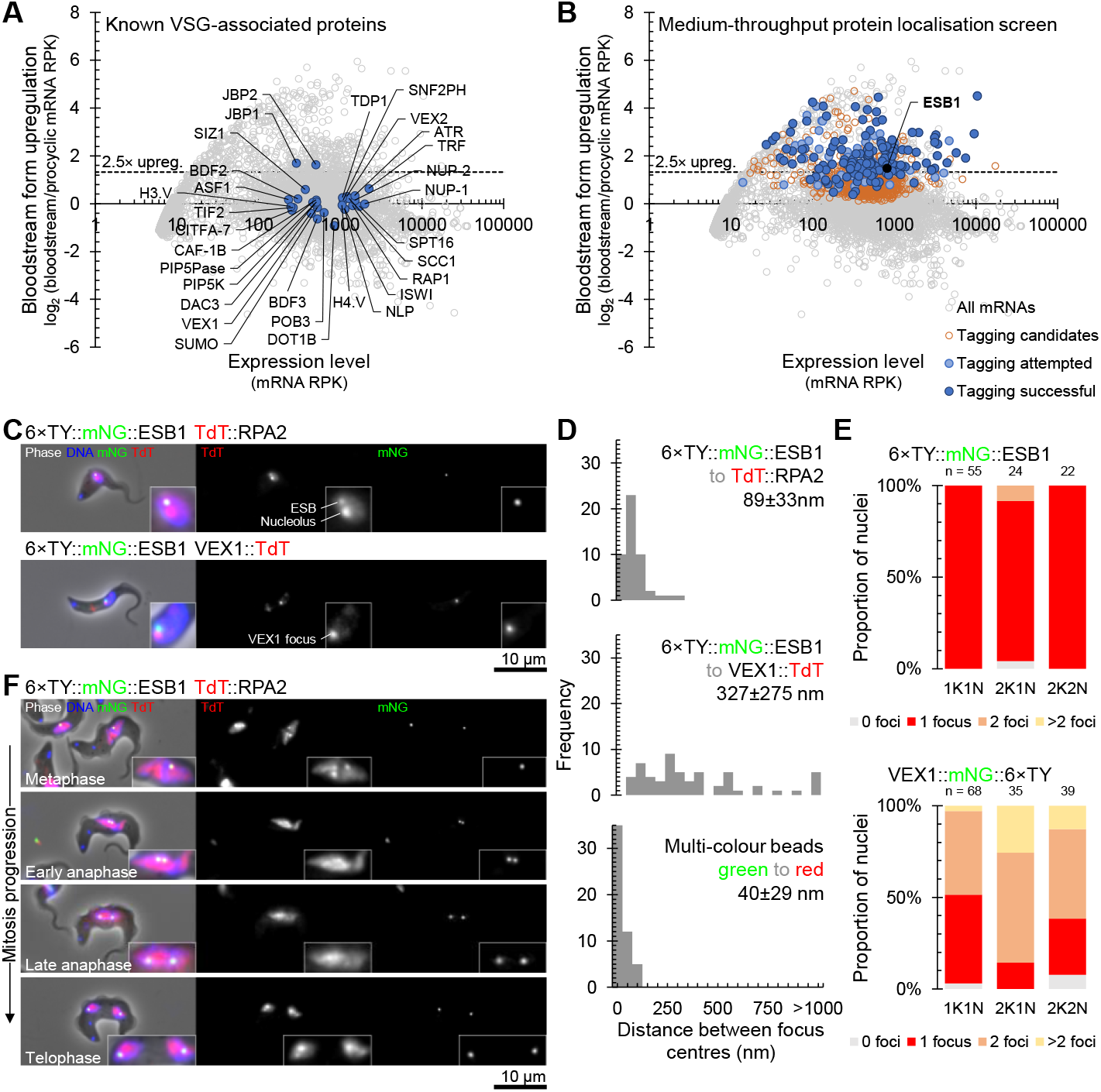
A protein localisation screen identified ESB1, the first ESB-specific protein. **A**,**B)** Degree of upregulation of *T. brucei* mRNAs in bloodstream forms previously determined by RNAseq ^43^ highlighting **A)** known VSG monoallelic expression-associated factors and **B)** candidates for tagging we selected, attempted and successfully localised. **C-F)** Microscopy analysis of ESB1 localisation. **C**,**D)** Localisation of mNG-tagged ESB1 relative to TdT-tagged RPA2 or VEX1 showing **C)** fluorescence microscope images of example G1 (1K1N) cells and **D)** distance between the ESB1 focus and either the RPA2 ESB focus or the nearest VEX1 focus. Multi-colour beads are a perfect colocalisation control. **E)** Number of mNG-tagged ESB1 or mNG-tagged VEX1 foci per nucleus in different cell cycle stages and **F)** fluorescence microscope images of mNG-tagged ESB1 foci in mitotic nuclei.

We performed a candidate protein localisation screen of proteins upregulated in the bloodstream form^28^ of unknown function, and identified the first ESB-specific protein. G1 bloodstream form nuclei have one extranucleolar ESB^3,29^. From 207 candidates only one (Fig. 1B), Tb427.10.3800, exhibited this localisation (Fig. 1C, Extended Data Fig. 2) whilst endogenous tagging in the procyclic form gave no detectable signal (Extended Data Fig. 2A). We named this protein ESB1.

We used well-characterised ESB markers to confirm the ESB1 localisation. Pol I is the founding component of the ESB and localises to both the nucleolus and the ESB in bloodstream forms^3^. ESB1 colocalised with Pol I (RPA2) at the ESB (Fig. 1C) confirmed by measurement of the distance between signal centre points (Fig. 1D). The ESB also has a VEX sub-complex involved in exclusion of inactive ESs^23^. ESB1 did not precisely colocalise with VEX1 (Fig. 1C,D). After S phase, cells still exhibit a single ESB before the nucleus undergoes closed mitosis – a second ESB only forms during anaphase^29^. ESB1 always localises to a single focus per nucleus (Fig. 1E) and only duplicates during mitosis (Fig. 1F). ESB1 is therefore specific to the ESB both spatially (localisation) and temporally (life cycle stage-specific expression and cell cycle-dependent localisation).

To determine ESB1 function, we generated a bloodstream form ESB1 conditional knockout (cKO) cell line (Extended Data Fig. 3A-E). ESB1 cKO gave undetectable levels of ESB1 by 24 h (Fig. 2C, Extended Data Fig. 3A) which caused a profound proliferation defect due to failure of cytokinesis and further rounds of organelle duplication (Fig. 2A,B).

**Fig. 2.**
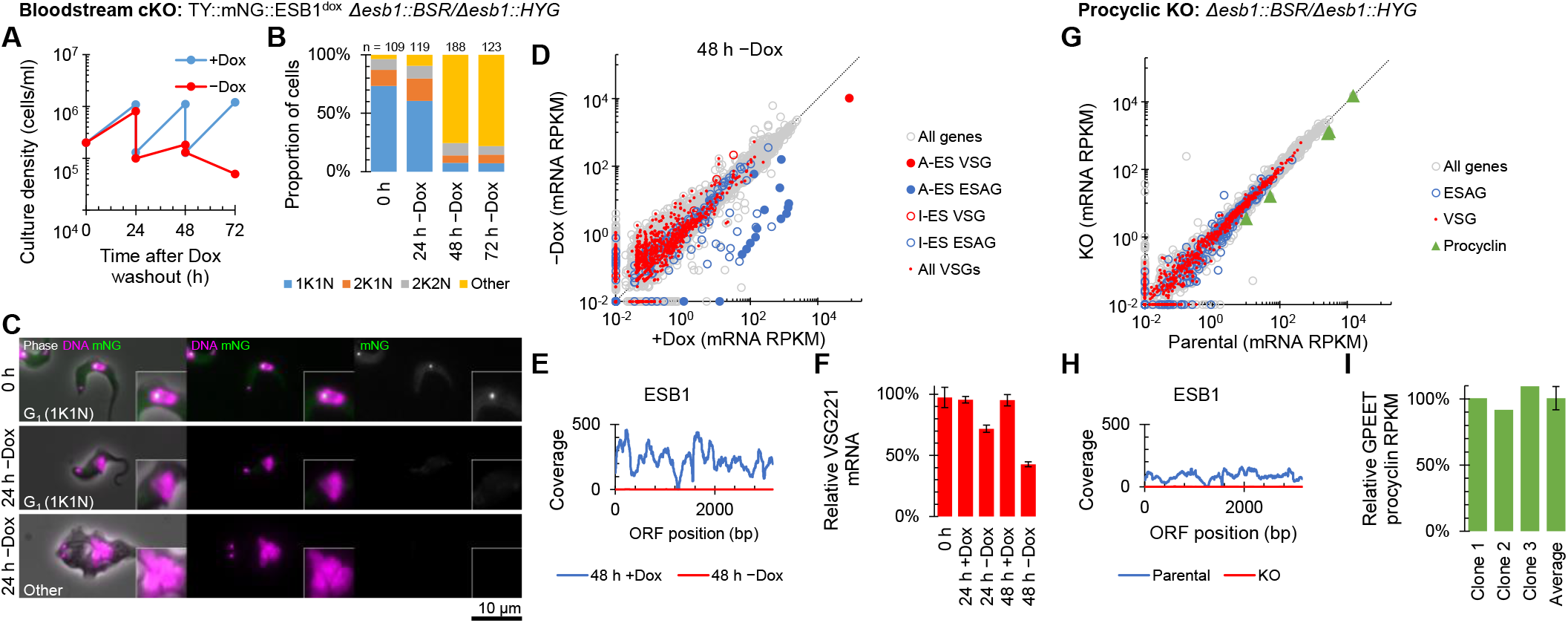
ESB1 is vital for bloodstream forms and required for transcription from the active VSG expression site. **A-F)** Cellular phenotype of bloodstream form ESB1 cKO. Exogenous mNG-tagged ESB1 expression was maintained with 10 ng/ml doxycycline (+Dox) in the bloodstream form ESB1 cKO (cell line validation in Extended Data Fig. 3A-D) and doxycycline washout (−Dox) induced the cKO phenotype. **A)** Culture growth (with subculture) and **B)** counts of morphologically abnormal (‘Other’) cells following washout. **C)** Example fluorescence microscope images showing mNG-tagged ESB1 signal before and 24 h after washout. **D)** Changes in mRNA abundance and **E)** ESB1 ORF read coverage determined by RNAseq 48 h after washout (full time series in Extended Data Fig. 3D). Active BES (A-BES) and inactive BES (I-BES) VSG and ESAG mRNAs along with other VSG mRNAs are highlighted in **D). F)** qRT-PCR quantitation of the A-ES VSG mRNA (VSG 427-2) after washout, *n* = 3. **G-I)** mRNA abundance phenotype of procyclic form ESB1 KO. **G)** Changes in mRNA abundance (further clonal cell lines in Extended Data Fig. 3E), **H)** ESB1 ORF read coverage and **I)** abundance of GPEET procyclin relative to the parental cell line determined by RNAseq.

To detect any effect on BES transcription, we used RNAseq to profile mRNA levels and showed ESB1 cKO caused a dramatic decrease (∼250×) in ESAG mRNAs. Mapping ESAGs to BESs^8^ showed that those with reduced mRNA levels were predominantly associated with the active BES (Fig. 2D, Extended Data Fig. 3E). The VSG in the active BES decreased ∼8× (Fig. 2D) which we confirmed by qPCR (Fig. 2F). The difference in these levels is expected given VSG mRNA has a longer half-life than ESAG mRNAs^30^. ESB1 cKO therefore reduces active BES transcription.

This cKO phenotype was recapitulated with the more experimentally amenable RNAi knockdown of ESB1 (Extended Data Fig. 4). The strength of the phenotype naturally led to appearance of RNAi escape sub-populations^31^; therefore, we analysed only early RNAi time points.

Procyclic forms lack an active BES, an ESB and do not express ESB1 although they use Pol I for expression of their surface coat protein (procyclin). We tested ESB1 cryptic function in procyclic forms by deleting both ESB1 alleles, which gave no apparent growth or morphology defect. RNAseq confirmed normal high expression of GPEET procyclin and no major changes to other mRNA transcripts (Fig. 2G-I, Extended Data Fig. 3F). ESB1 is therefore vital in bloodstream forms for monoallelic VSG expression but not required in procyclic forms.

We next determined whether ESB1 is required for the normal molecular composition of the ESB. We generated a panel of cell lines carrying the inducible ESB1 RNAi construct and tagged ESB proteins: RPA2, SUMO (as the ESB is associated with a highly SUMOylated focus^21^), VEX1 or VEX2 (Fig. 3A-H). As shown by others, the ESB focus of RPA2 was visible in 40% of G1 nuclei (ie when not occluded by nucleolar RPA2)^3^ and the highly SUMOylated focus in ∼60% of G1 nuclei^21^. After 24 h induction of ESB1 RNAi, RPA2 and SUMO were more dispersed through the nucleus and fewer nuclei had an ESB focus in both morphologically normal and abnormal cells (Fig. 3A-D). As seen previously, VEX1 and VEX2 localised to 1 or 2 foci in the nucleus. After 24 h induction of ESB1 RNAi the localisation pattern was unchanged, both in morphologically normal and abnormal cells (Fig. 3E-H). This indicates ESB1 is necessary for both recruitment of Pol I to the ESB and higher local SUMOylation to form the HSF, but not formation of VEX foci.

**Fig. 3.**
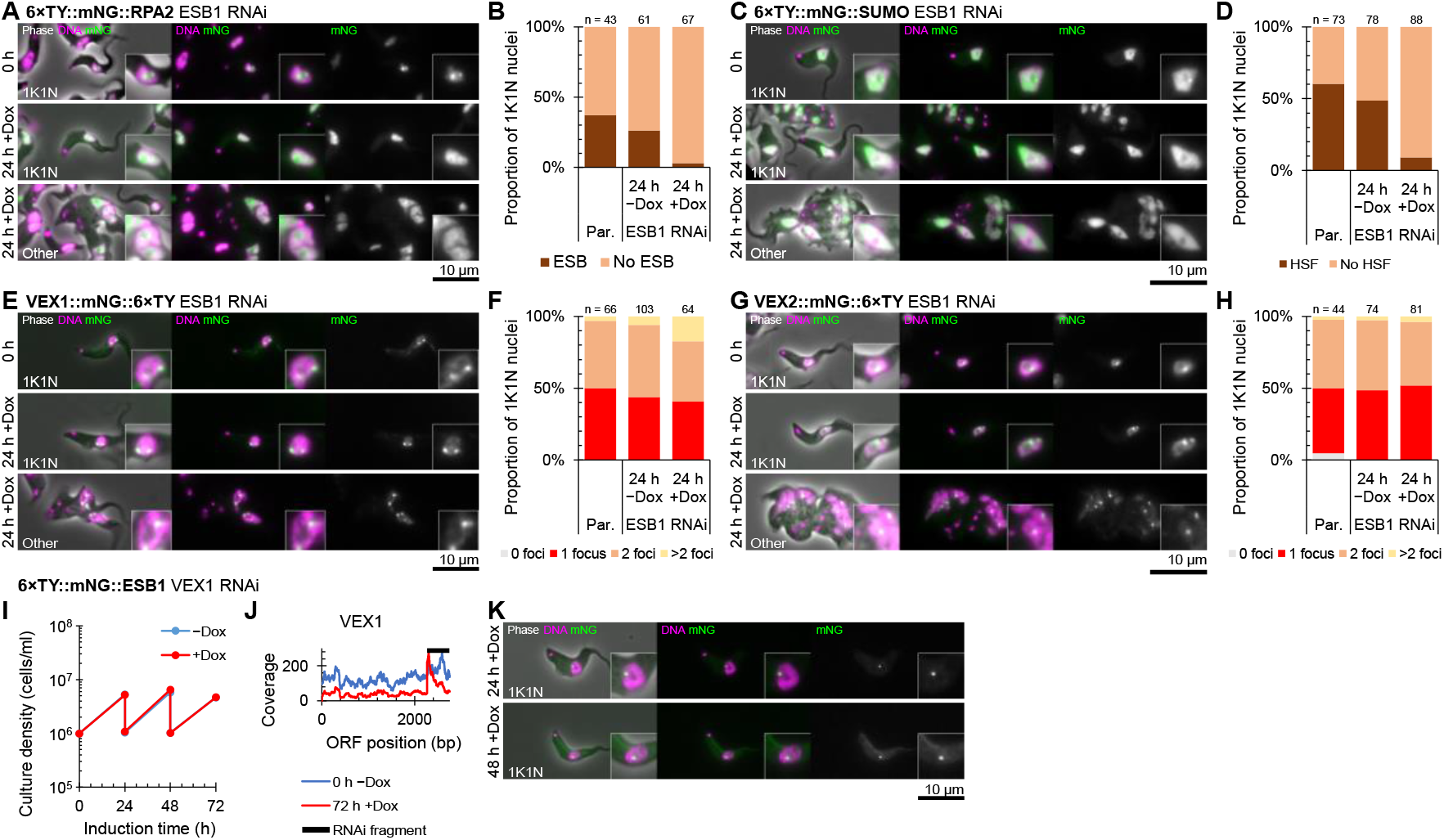
ESB1 is required for formation of a subset of ESB substructures. **A-H)** Effect of doxycycline-inducible ESB1 RNAi knockdown (knockdown characterised in Extended Data Fig. 4) on mNG-tagged **A-B)** RPA2, **C-D)** SUMO, **E-F)** VEX1 and **G-H)** VEX2 localisation. Each cell line was maintained without doxycycline (−Dox) then induced with 1 mg/ml doxycycline (+Dox). **A**,**C**,**E**,**G)** Example fluorescence microscope images show tagged protein localisation before induction and in morphologically normal and abnormal cells after 24 h RNAi induction. **B**,**D**,**F**,**H)** Counts of the number of cells with an **B)** RPA2-containing ESB focus or **D)** highly SUMOylated focus (HSF) and the number of **F)** VEX1 or **H)** VEX2 in comparison to the parental (“Par.”, no RNAi) cell line. **I-K)** Effect of doxycycline-inducible VEX1 RNAi knockdown on mNG-tagged ESB1 localisation. **I)** Culture growth (with subculture) and **J)** VEX1 ORF read coverage showing effective knockdown determined by RNAseq. **K)** Example fluorescence microscope images showing mNG-tagged ESB1 localisation after VEX1 knockdown.

The inverse, whether the ESB1 focus is VEX1-dependent, was analysed by depleting VEX1 using RNAi and observing tagged ESB1 (Fig. 3I-K). RNAseq profiling of mRNA confirmed VEX1 knockdown with no growth defect (Fig. 3I,J) and, as previously described^23^, derepression of inactive BESs (data not shown). Tagged ESB1 localisation was unchanged on VEX1 knockdown (Fig. 3K); therefore, formation of a singular ESB is not dependent on VEX1 repression of inactive BESs.

We then asked whether ESB1 overexpression could force ectopic BES expression and/or supernumerary ESB formation. Significant overexpression was achieved using a cell line with an additional inducible tagged ESB1 locus (using 100 ng/ml doxycycline) (Fig. 4A-F, Extended Data Fig. 3B). In contrast to ESB1 cKO, overexpression gave a small growth reduction and some cytokinesis defects (Fig. 4A,B). Overexpressed ESB1 still localised to the ESB, although with more dispersion in the nucleoplasm and cytoplasm, for both morphologically normal and abnormal cells (Fig. 4C). ESB1 overexpression in a cell line also expressing tagged RPA2 showed Pol I was not dispersed and still localised at the single nucleolus and ESB (Fig. 4G-I). In bloodstream forms ESB1 overexpression therefore does not alter ESB number or form.

**Fig. 4.**
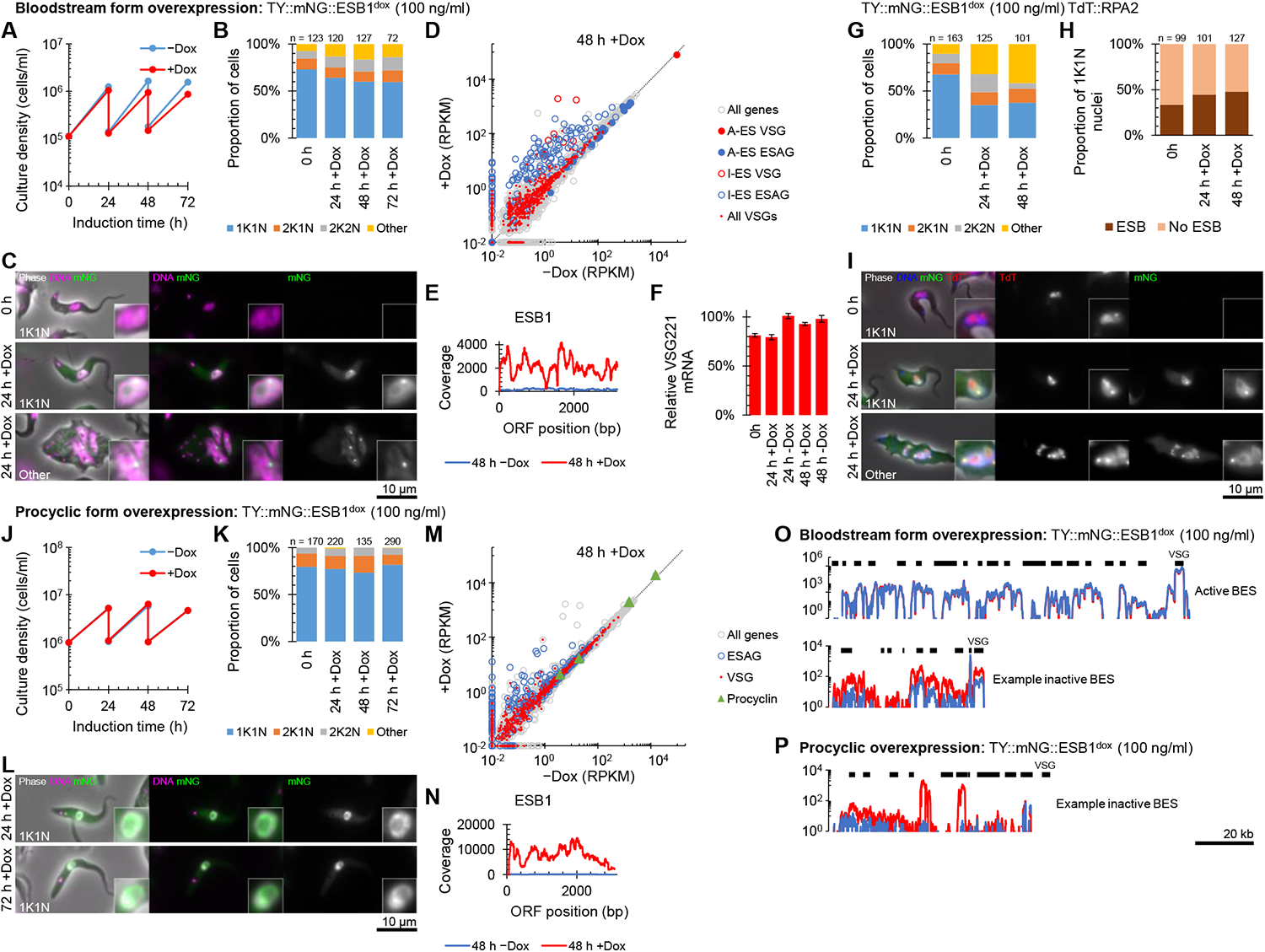
ESB1 overexpression activates inactive BESs without affecting ESB formation. **A-F)** Cellular phenotype of mNG-tagged ESB1 overexpression in bloodstream forms induced with 100 ng/ml doxycycline (+Dox). **A)** Culture growth (with subculture) and **B)** counts of morphologically abnormal (‘Other’) after induction. **C)** Example fluorescence microscope images of mNG-tagged ESB1 signal in morphologically normal cells before and morphologically normal and abnormal cells 24 h after overexpression induction. **D)** Changes in mRNA abundance and **E)** ESB1 ORF read coverage determined by RNAseq 48 h after washout (full time series in Extended Data Fig. 5A). Active BES (A-BES) and inactive BES (I-BES) VSG and ESAG mRNAs along with other VSG mRNAs are highlighted in **D). F)** qRT-PCR quantitation of the A-ES VSG mRNA (VSG 427-2) after washout, *n* = 3. **G-I)** Effect of mNG-tagged ESB1 overexpression on TdT-tagged RPA2 localisation. Counts of **G)** morphologically abnormal cells, **H)** the number of cells with an RPA2-containing ESB focus and **I)** fluorescence microscope images show tagged protein localisation before induction and in morphologically normal and abnormal cells after ESB1 overexpression. **J-N)** Cellular phenotype of mNG-tagged ESB1 overexpression in procyclic forms induced with 100 ng/ml doxycycline. **J)** Culture growth (with subculture) and **K)** counts of morphologically abnormal (‘Other’) after induction. **L)** Example fluorescence microscope images of overexpressed mNG-tagged ESB1 in procyclic forms. **M)** Changes in mRNA abundance and **N)** ESB1 ORF read coverage determined by RNAseq 48 h after washout (full time series in Extended Data Fig. 5B). **O-P)** BES read coverage, determined by RNAseq 48 h after ESB1 overexpression, of **O)** the active and an example inactive BES in bloodstream forms and **P)** an example inactive BES in procyclic forms.

RNAseq transcriptome profiling of the ESB1 overexpression cell line showed a dramatic increase mRNA levels, up to ∼100×, for VSGs and ESAGs in inactive BESs while the mRNA level of VSG and ESAGs in the active BES remained unchanged (Fig. 4D, Extended Data Fig. 5A and confirmed using qRT-PCR in Fig. 4F) arising from a ∼10× increase in ESB1 mRNA (Fig. 4E). ESB1 overexpression is therefore sufficient to cause activation of inactive VSG BESs whilst expression of procyclic form-specific surface proteins remained low (Extended Data Fig. 5A).

Finally, we forced expression of tagged ESB1 in procyclic form cells (Fig. 4J-N, Fig. S5B). Significant expression produced no growth or cytokinesis defects (Fig. 4J,K). Tagged ESB1 was nuclear but did not localise to a single extranucleolar ESB-like focus (Fig. 4L). RNAseq analysis showed a large increase (up to ∼200×) in mRNA level for ESAGs, consistent with transcription initiation at BES promoters which are normally inactive in the procyclic form (Fig. 4M). In this particular strain, we interrogated expression of the ESAGs and VSG from the sequenced BES^32^. Every ESAG in this BES was upregulated, typically ∼3-5× and up to ∼80× (Fig. 4O,P). In contrast, VSG mRNAs (both published and from our *de novo* assembly of the transcriptome) were not strongly upregulated (Fig. 4M). We did not see transcript from VSG 10.1, found in the sequenced BES, nor upregulation of any of the VSGs commonly expressed by this strain in bloodstream forms during mouse infection^33^. This is despite ∼50× overexpression of ESB1 transcript relative to endogenous bloodstream form expression (Fig. 4E, Fig. 4N). As for tagged ESB1 overexpression in the bloodstream form, procyclin mRNA levels also remained unchanged (Fig. 4M).

Hence ESB1 expression in procyclic forms activates BES transcription without forming an ESB; however, transcription is either not fully processive to the most distal gene (VSG) or there is additional machinery required for VSG transcript maturation, processing and/or stability not expressed in the procyclic form.

Antigenic variation in *T. brucei* relies on monoallelic expression of the VSG gene in the active BES. The transcriptional activation of the active BES is counterbalanced by a strong repression of all other BESs. Our work has provided insights into ESB subdomains which orchestrate these different functions.

We have identified ESB1 as the first ESB-specific protein and as a BES transcription activator. We show that ESB1 is necessary for transcription from the active BES and is specific to only VSG-containing ESs (BESs and MESs – ESs containing metacyclic VSGs expressed in the mammalian infective parasite in the insect) and not procyclin loci. The predicted RING domain of ESB1 has weak similarity to ubiquitin E3 ligases, suggesting protein rather than DNA interaction. Importantly, ectopic expression of ESB1 in the procyclic form was sufficient to upregulate ESAGs located within a BES. However, ESB1 alone in procyclic forms is not sufficient for fully processive BES transcription and/or VSG mRNA processing.

ESB1 alone was not sufficient to support formation of the Pol I and ESB1 focus, as overexpression of ESB1 did not give rise to an ESB-like body/bodies in the procyclic form or supernumerary ESB-like bodies in the bloodstream form. Moreover, multiple active BESs in multiple ESBs is not a stable state: in cells forced to express two VSGs from two BESs, both were recruited to a single ESB^34^. Given this and our ESB1 overexpression results suggests that the reasons for ESB absence (procyclic forms) or singularity (bloodstream forms) are likely to be more complex than a threshold level of ESB1 protein. Singularity achieved by deterministic or emergent properties (such as Otswald ripening if the compartment is formed by phase separation) may also have important implications for switching between BESs.

Our work, taken with that of others, shows that the ESB is a complex nuclear body with multiple subdomains. The defining subdomain is a focus of Pol I around the active BES^3^, which also contains basal Pol I transcription factors^35^ and ESB1. This sits within a highly SUMOylated focus (HSF)^21^. ESB1 is required for assembly of this subdomain. The BES is found in close proximity to one of the spliced leader array alleles^25^. Pol II transcription of this array generates spliced leader RNA necessary for processing of all transcripts into mRNA. Each spliced leader array allele is found in a Pol II transcription focus^27^ and the proximity of one to the ESB BES/Pol I subdomain provides a mechanism for efficient processing of the large amount of VSG mRNA. BES association with the ESB spliced leader array/Pol II subdomain requires VEX2^25^ and the ESB BES/Pol I subdomain overlaps or is adjacent to one VEX1 and VEX2 nuclear focus^23–25^. We show that assembly of these foci are separable, with assembly of the VEX foci not dependent on ESB1 and vice-versa. Importantly we show that the Pol I and ESB1 focus is strictly singular. A new appreciation of the ESB in terms of spatially defined subdomains raises the possibility that this reflects an intrinsic functional architecture.

## Methods

### Parasite strains and cell culture

*Trypanosoma brucei brucei* strain Lister 427 was selected for monomorphic bloodstream form (BSF) experiments because its expression sites have been cloned and sequenced^36^, and are assembled into contigs in the 2018 re-sequence^8^. BSFs were grown in HMI-9^37^ at 37°C with 5% CO_2_, maintained under approximately 2×10^6^ cells/ml by regular subculture. We confirmed that the active BES was BES1 containing VSG 427-2 (see Transcriptomic analysis).

*T. brucei brucei* strain TREU927 was selected for procyclic form (PCF) experiments as it is the original genome strain^38^ with a well-annotated genome and was used for our genome-wide protein localisation project in PCFs^28,39^. PCFs were grown in SDM-79^40^ at 28°C, maintained between approximately 6×10^5^ and 2×10^7^ cells/ml by regular subculture.

To enable CRISPR/Cas9-assisted genetic modifications and to use doxycycline-inducible genetic modifications we used PCF and BSF cell lines which expresses T7 polymerase, Tet repressor, Cas9 and PURO drug selectable marker. These cell lines were generated using pJ1339, an expression construct which carries homology arms for integration in the tubulin locus. We have previously described the TREU927 PCF 1339 cell line^41^. To generate the Lister 427 BSF 1339 cell line, pJ1339 was linearised by restriction digest with *Hind*III and transfection into BSFs (see Electroporation and drug selection).

Culture density was measured with a haemocytometer (BSFs) or a CASY model TT cell counter (Roche Diagnostics) with a 60 µm capillary and exclusion of particles with a pseudo diameter below 2.0 µm (PCFs).

### Electroporation and drug selection

Electroporation was used to transfect *T. brucei* with linear DNA constructs which have 5′ and 3′ homology arms, leading to construct integration into the target locus by homologous recombination. 1 to 5 µg of linearised plasmid DNA or DNA from the necessary PCR reactions was purified by either phenol chloroform extraction (for the medium-throughput bloodstream form localisation screen) or ethanol precipitation (other experiments) then mixed with approximately 3×10^7^ cells for BSFs or 1×10^7^ cells for PCFs resuspended in 100 µl of Tb-BSF buffer^42^. Transfection was carried out using program X-001 of the Amaxa Nucleofector IIb (Lonza) electroporator in 2 mm gap cuvettes. Following electroporation, cells were transferred to 10 ml pre-warmed HMI-9 (BSFs) or 10 ml SDM-79 (PCFs) for 6 h then the necessary selection drugs were added to select for cells with successful construct integration. Clonal cell lines were generated for all cell lines, except those from medium throughput tagging, by limiting dilution cloning.

All cultures were maintained with periodic drug selection for any genetic modifications, using the necessary combination of 0.2 µg/ml (BSF) or 1.0 µg/ml (PCF) Puromycin Dihydrochloride, 5 µg/ml (BSF) or 10 µg/ml (PCF) Blasticidin S Hydrochloride, 2.0 µg/ml (BSF) or 15 µg/ml (PCF) G-418 Disulfate, 5 µg/ml (BSF) or 25 µg/ml (PCF) Hygromycin B, 2.5 µm/ml (BSF) or 5 µm/ml Phleomycin. Cultures were maintained without drug selection for at least one subculture prior to an experiment.

### Medium-throughput BSF localisation screen for ESB proteins

Our localisation-based screen for ESB proteins used endogenous tagging with a fluorescent protein (see Endogenous tagging). Tagging candidates were selected based on previously published mRNA abundance data determined by RNAseq^43^ by selecting genes with transcripts significantly upregulated (*p* < 0.05, Student’s T test) in the BSF relative to the PCF and prioritising those >2.5× upregulated (Fig. 1B). We further prioritised genes with unknown function, and excluded VSG genes and pseudogenes, ESAGs, genes related to ESAGs (GRESAGs) and known invariant surface glycoproteins (ISGs). Some known proteins were, however, tagged as controls, such as ISG65 and GPI-PLC. We also used other transcriptomic and ribosome footprinting datasets for further manual prioritisation^43–46,46,47^.

Tagging was primarily at the N terminus. The C terminus was tagged if the protein had a predicted signal peptide. We attempted tagging of 207 proteins with a 73% success rate generating a 153 tagged cell lines. Of these, 7 had a nuclear signal (0).

### Endogenous tagging

N or C terminal tagging by modification of gene endogenous loci was carried out as previously described, using long primer PCR to generate tagging constructs and, for BSF tagging, PCR to generate DNA encoding sgRNA with a T7 promoter^48,49^. Primer design, PCR using the pPOT series of plasmids as templates and sgDNA production were carried out as previously described^48,49^. mNeonGreen (mNG)^50^ was used for all tagging with a green fluorescent protein. pPOT v4 Blast mNG was used for the medium-throughput bloodstream form localisation screen. pPOT v6 Blast 6×TY::mNG was used for other experiments. In this construct 3×TY epitope tags lie either side of the mNG (3×TY::mNG:: 3×TY), however for simplicity we refer to this as a 6×TY::mNG tag. tdTomato (TdT) and pPOT v4 Hyg TdT was used for tagging with a red fluorescent protein for colocalisation experiments.

For ESB1 tagging in PCFs, where no fluorescent signal from tagging was detected, we generated a clonal cell line by limiting dilution and confirmed the correct fusion of the mNG CDS to the ESB1 CDS was achieved (see Endogenous locus ORF modification/loss PCR validation) (Extended Data Fig. 2A,B).

For ESB1 tagging in BSFs, to test whether the fluorescent protein tag on ESB1 caused a protein localisation or function defect we confirmed that the C terminally-tagged protein had the same localisation as the N terminally-tagged (Extended Data Fig. 2C,D). *T. brucei* are diploid.

Therefore we also confirmed, by deletion of the untagged allele, that expression of N terminally-tagged ESB1 in the absence of the wild-type allele gave the same localisation (Extended Data Fig. 2E,F) and we saw no morphological or cell growth defect (data not shown).

### Gene knockout and conditional knockout

Gene knockout was carried out using long primer PCR to generate deletion and sgRNA constructs^48,49^. As for endogenous tagging, this was carried out as previously described^48,49^. pPOT v7 Hyg and pPOT v6 Blast were used for generation of deletion constructs. We confirmed knockout by testing for loss of the target gene CDS and replacement of the target gene CDS with the drug selection marker (see Endogenous locus ORF modification/loss PCR validation). We were unable to generate a BSF ESB1 deletion cell line and therefore used a conditional knockout (cKO) approach by first generating a cell line capable of doxycycline-inducible exogenous ESB1 expression.

For exogenous ESB1 expression or overexpression, the Tb927.10.3800 ORF was amplified by PCR from *T. b. brucei* TREU927 gDNA and cloned into variants of pDex577^51^ and pDex777^52^ with an 1×TY::mNG combined fluorescence reporter and epitope tag. These are doxycycline-inducible constructs using a T7 promoter for the inducible expression which carry homology arms for integration into the transcriptionally silent minichromosome repeats. Each pDex577/pDex777 construct was linearised by restriction digest with *Not*I and transfected into BSFs and PCFs (see Electroporation and drug selection).

We titrated doxycycline concentrations to select concentrations which achieved desirable tagged ESB1 expression levels in BSFs and PCFs, using a combination of light microscopy (to evaluate fluorescence intensity of mNG relative to cell lines with an endogenous ESB1 mNG tag), Western blots (see Western blotting) (Extended Data Fig. 3A,B) and RNAseq (see Transcriptomic analysis). We selected induction conditions to give (i) expression comparable to endogenous expression levels in BSFs (pDex577 with 10 ng/ml doxycycline Extended Data Fig. 3B), (ii) overexpression sufficient to generate an aberrant BSF phenotype (pDex577 with 100 ng/ml doxycycline, Extended Data Fig. 3B) or (iii) high overexpression in PCFs (pDex777 with 1 mg/ml doxycycline, Fig. 4N).

RNAseq confirmed no major perturbation of cellular transcripts in BSFs expressing tagged ESB1 using pDex577 with 10 ng/ml doxycycline (Extended Data Fig. 3C). We then deleted both endogenous ESB1 alleles (Extended Data Fig. 3D) while maintaining the cell line with 10 ng/ml doxycycline to generate the cKO cell line. The cKO phenotype was observed by washing doxycycline out of the culture (see Induction time series).

### Endogenous locus ORF modification/loss PCR validation

Key endogenous locus modifications were validated using PCR using genomic DNA (gDNA) extracted from the modified cell line using the DNeasy Blood & Tissue Kit (Qiagen) as the template. Primer pairs were designed spanning from the endogenous DNA sequence to the DNA introduced by homologous recombination. For deletions, the gene 5′ UTR to the drug selection marker ORF (Extended Data Fig. 2F, Extended Data Fig. 3D), and for tagging, the gene ORF to the fluorescent tag ORF (Extended Data Fig. 2B,F). PCR product size was checked by agarose gel electrophoresis. Primer sequences used are listed in Extended Data Table 2. For genetic modifications where both gene alleles were modified (e.g. gene deletions) the first allele was modified, the modification confirmed by PCR using gDNA as the template, then the second allele was modified.

### Inducible RNAi knockdown

For inducible ESB1 or VEX1 RNAi knockdown, we amplified a fragment of the target genes’ ORFs and cloned them into a new doxycycline-inducible RNAi construct, pDRv0.5. The resulting plasmid had two copies of the amplicon cloned in reverse complement separated by a 150 nt stuffer fragment, with transcription of the resulting “stem-loop” driven by two T7 promoters under the control of doxycycline that flanked the insert (see Extended Data Table 3 for primer sequences). Cell were transfected with *Not*I linearised plasmid and transgenic cells selected using Hygromycin B (see Electroporation and drug selection). RNAi knockdown was induced using 1 µg/mL doxycycline.

To confirm these RNAi constructs gave effective knockdown we introduced them into cell lines expressing an endogenously tagged copy of the target protein whose knockdown could be confirmed by light microscopy (Extended Data Fig. 4C) and/or Western blot (Extended Data Fig. 4D) and/or RNAseq to determine transcript abundance of the target gene (see Transcriptomic analysis) (Fig. 3J).

For ESB1 RNAi knockdown in cell lines expressing endogenously tagged RPA2, SUMO, VEX1 or VEX2 we confirmed effective ESB1 knockdown, and confirmed that that escape from ESB1 RNAi had not occurred, by checking for the expected growth rate defect and determining the proportion of cells at different cell cycle stages in the population (see Induction time series).

### Western blotting

We used Western blotting to confirm molecular weight and expression level of endogenously tagged and exogenously (over)expressed proteins. An anti-mNG (mouse monoclonal IgG2c, ChromoTek 32f6, RRID:AB_2827566) or anti-TY (from BB2 hybridoma, mouse monoclonal IgG1^53^) primary antibody and anti-mouse HRP-conjugated secondary antibody were used using standard protocols.

### Induction time series

Doxycycline-inducible and cKO cell lines were analysed as induction time series with paired induced and uninduced samples. Cells in logarithmic growth were subcultured to 1×10^5^ cell/ml (BSFs) or 1×10^6^ cells/ml (PCFs), one sample without and one with the necessary concentration of doxycycline for induction. Each 24 h, the culture density was measured, samples taken, then the remaining cells subcultured to 1×10^5^ cell/ml (BSFs) or 1×10^6^ cells/ml (PCFs), including doxycycline in the induced sample. For cultures with a strong growth defect, the culture was centrifuged at 1200 g for 5 min then the cell pellet was resuspended in fresh medium, and doxycycline added if needed, in order to maintain constant conditions. Each set of samples included a sample for light microscopy (to analyse cell morphology and cell cycle progression, see Microscopy) and any RNA, protein, or other samples.

### Microscopy

Light microscopy was carried out on live cells adhered to glass with DNA stained with Hoechst 33342, as previously described^54^. Images were captured on a DM5500 B (Leica) upright widefield epifluorescence microscope using a plan apo NA/1.4 63× phase contrast oil-immersion objective (Leica, 15506351) and a Neo v5.5 (Andor) sCMOS camera using MicroManager^55^.

Cell cycle progression was evaluated from microscope images. In the normal *T. brucei* cell cycle kinetoplast (K, mitochondrial DNA) division precedes nuclear (N) division, giving 1K1N, 2K1N then 2K2N cells prior to cytokinesis. These approximately correspond to G_1_, S and post-mitosis to cytokinesis phases respectively. K/N number in cells were manually counted as a measure of cell cycle progression, with abnormal K/N numbers classified as ‘other’

Spacing of point-like structures, one in the green channel and one in the red, was carried out by fitting a Gaussian in the X and Y directions to determine the signal centre point in each channel then calculating their separation, as previously described^56^. Prior to analysis, images were corrected for chromatic aberration as previously described^57^ using a sample of 0.1 µm TetraSpeck multi-colour fluorescent beads (ThermoFisher) adhered to glass as a reference sample. Measurement error was determined by measuring green-red spacing using the multi-colour fluorescent beads.

For blinded counts of nuclear structures, one researcher identified and cropped in-focus nuclei (based on Hoechst 33342 signal) from 1K1N cells in microscopy fields of view from a mixture of test and control samples. Each cropped image was saved with a randomised unique file name, recording the source sample in an index file. A second researcher then classified the image files containing single nuclei, then the analysis was unblinded using the index file.

### Transcriptomic analysis

Total RNA samples from 10^8^ cells were extracted using the RNeasy Mini Kit (Qiagen). For RNAseq, mRNA was enriched by polyA selection, cDNA generated by reverse transcription using a poly-dT primer then subject to 100 bp paired end sequencing with a nominal insert size of 200 bp and >70,000,00 reads per sample.

To quantify transcript abundance, fastq reads were mapped to the appropriate predicted transcriptome using BWA-MEM (version 0.7.17) with default settings. TriTrypDB^58,59^ release 45 of the *T. brucei* Lister 427 2018 annotated transcripts for 427 was used for BSF samples and *T. brucei* TREU927 annotated transcripts for 927 PCF samples. An additional contig for the single sequenced and assembled *T. brucei* TREU927 BES^32^ was generated by combining NCBI GenBank accession numbers AC087700 (BES) and AF335471 (VSG), from which the ESAG and VSG open reading frames were identified and appended to the *T. brucei* TREU927 predicted transcriptome. To quantify uniquely mapped read coverage the resulting bam file was filtered to include reads mapped in proper pairs and exclude reads not mapped, exclude read secondary alignments and exclude read PCR or optical duplicates with a MAPQ>10, using samtools view (version 1.7) with flags -q 10, -F 0×504 and -f 0×02. RPKM was calculated from the output of samtools (version 1.7) idxstats. Mean coverage was calculated from the output of samtools depth with flags -aa and -d 10000000.

To confirm changes to active BES VSG expression, we carried out quantitative reverse transcription PCR (qRT-PCR) using a one-step protocol directly from total RNA using primers specific to VSG 427-2 and β-tubulin (Extended Data Table 4). RNA samples were diluted to approximately equal concentration based on OD_260_. A dilution series of six three-fold steps from 1:3^0^ (1:1) to 1:3^6^ (1:279) dilutions of RNA from the parental cell line was used to confirm critical cycles for both VSG 427-2 and tubulin fell in the linear range. Mean VSG 427-2 and tubulin critical cycle was determined in triplicate using 1:10 diluted RNA samples. Samples were compared using VSG 427-2 to tubulin critical cycle ratio.

For *de novo* transcriptome assembly we used Trinity (version 2.11.0) guided by Harvard FAS best practices^60^. Sequencing errors were first corrected using Rcorrector (version 1.0.4)^61^ and uncorrectable reads were removed, then any remaining adapters and low quality read sections trimmed with Trim Galore! (version 0.6.0) with flags --length 36, -q 5, --stringency 1 and -e 0.1. Finally, sequences at the end of the read which exactly matched 4 or more bases of the 3′ end of the *T. brucei* spliced leader sequence were trimmed using a custom python script. Trinity was then used using default settings to generate the *de novo* assembly.

## Acknowledgements

This work was supported by the Wellcome Trust [104627/Z/14/Z, 108445/Z/15/Z, 211075/Z/18/Z], JS is supported by a David Fell Fellowship and RJW is a Sir Henry Dale Fellow.

## Author contributions

Conceptualization, KG, JDS, RJS, SD; Formal Analysis, RJW; Funding, KG, RJW, SD, JDS; Investigation, LLE, BH, CH, JDS, RJW, SD; Supervision, KG, JDS, RJS, SD; Visualization, RJW, LLE; Writing, KG, JDS, RJW, SD, LLE.

---

**Extended Data Table 1.** Previously described proteins involved in VSG expression or monoallelic exclusion.

| Group | Protein | Gene ID | Reference |
| --- | --- | --- | --- |
| ESB-associated VSG exclusion | VEX1 | Tb927.11.16920 | 23 |
|  | VEX2 | Tb927.11.13380 | 24 |
| ESB-associated transcription activators | SUMO | Tb927.5.3210 | 21 |
|  | SIZ1 | Tb927.9.11070 | 21 |
|  | SNF2PH | Tb927.3.2140 | 22 |
| Pol I basal transcription factors | CITFA-7 | Tb927.7.2600 | 35,62 |
| Cohesin | SCC1 | Tb927.7.6900 | 29 |
| Telomere-associated | ORC1 | Tb927.11.7216 | 9 |
|  | RAP1 | Tb927.11.370 | 12 |
|  | TIF2 | Tb927.3.1560 | 10,63 |
|  | TRF | Tb927.10.12850 | 10,64 |
|  | JBP1 | Tb927.11.13640 | 11 |
|  | JBP2 | Tb927.7.4650 | 11 |
| Specialised chromatin & remodelling | DOT1B | Tb927.1.570 | 15 |
|  | CAF-1B | Tb927.10.7050 | 13 |
|  | ASF1 | Tb927.1.630 | 13 |
|  | ISWI | Tb927.2.1810 | 16 |
|  | NLP | Tb927.2.1810 | 17 |
|  | BDF2 | Tb927.10.7420 | 18 |
|  | BDF3 | Tb927.11.10070 | 18 |
|  | POB3 | Tb927.10.14390 | 14 |
|  | SPT16 | Tb927.3.5620 | 14 |
|  | DAC3 | Tb927.2.2190 | 19 |
|  | SAP | Tb927.10.4440 | 65 |
| Nuclear envelope | NUP-1 | Tb927.2.4230 | 66 |
|  | NUP-2 | Tb927.9.6460 | 67 |
| Specialised chromatin | TDP1 | Tb927.3.3490 | 20,68 |
|  | H3.V | Tb927.10.15350 | 8 |
|  | H4.V | Tb927.2.2670 | 8 |
| Chromatin | H1 | Tb927.11.1790, Tb927.11.1800, Tb927.11.1860, Tb927.11.1870, Tb927.11.1880 | 69 |
|  | H3 | Tb927.1.2430, Tb927.1.2450, Tb927.1.2470, Tb927.1.2490, Tb927.1.2510, Tb927.1.2530, Tb927.1.2550 | 13 |
| DNA damage | ATR | Tb927.11.14680 | 70 |
| Membrane | PIP5Pase | Tb927.11.6270 | 71,72 |
|  | PIP5K | Tb927.10.3890 | 71 |
|  | PLC | Tb927.11.5970 | 71 |

**Extended Data Table 2.**
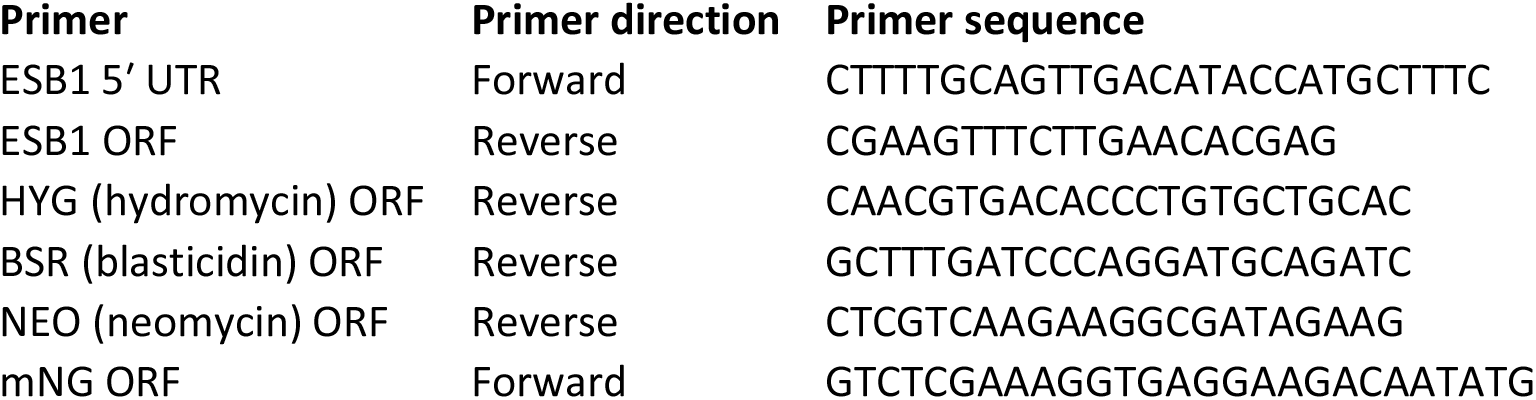
Primer sequences used for PCR validation of genetic modification.

**Extended Data Table 3.**
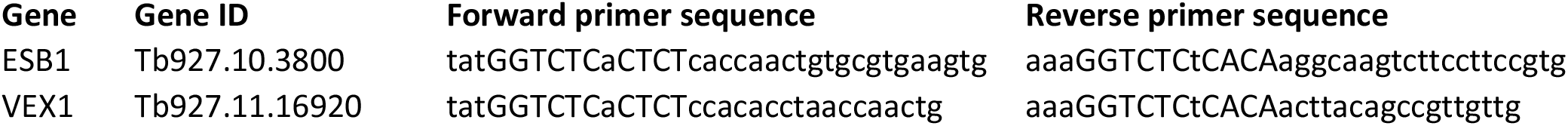
Primer sequences used for generation of RNAi constructs.

**Extended Data Table 4.**
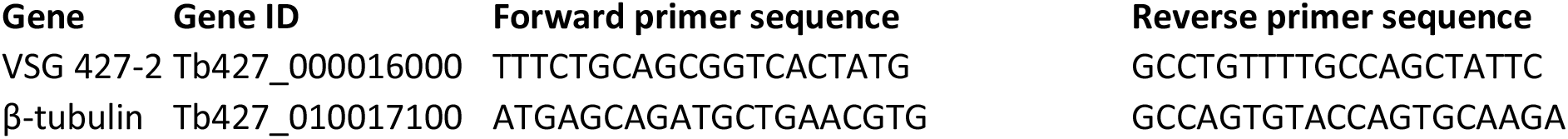
Primer sequences used for qRT-PCR.

**Extended data fig. 1.**
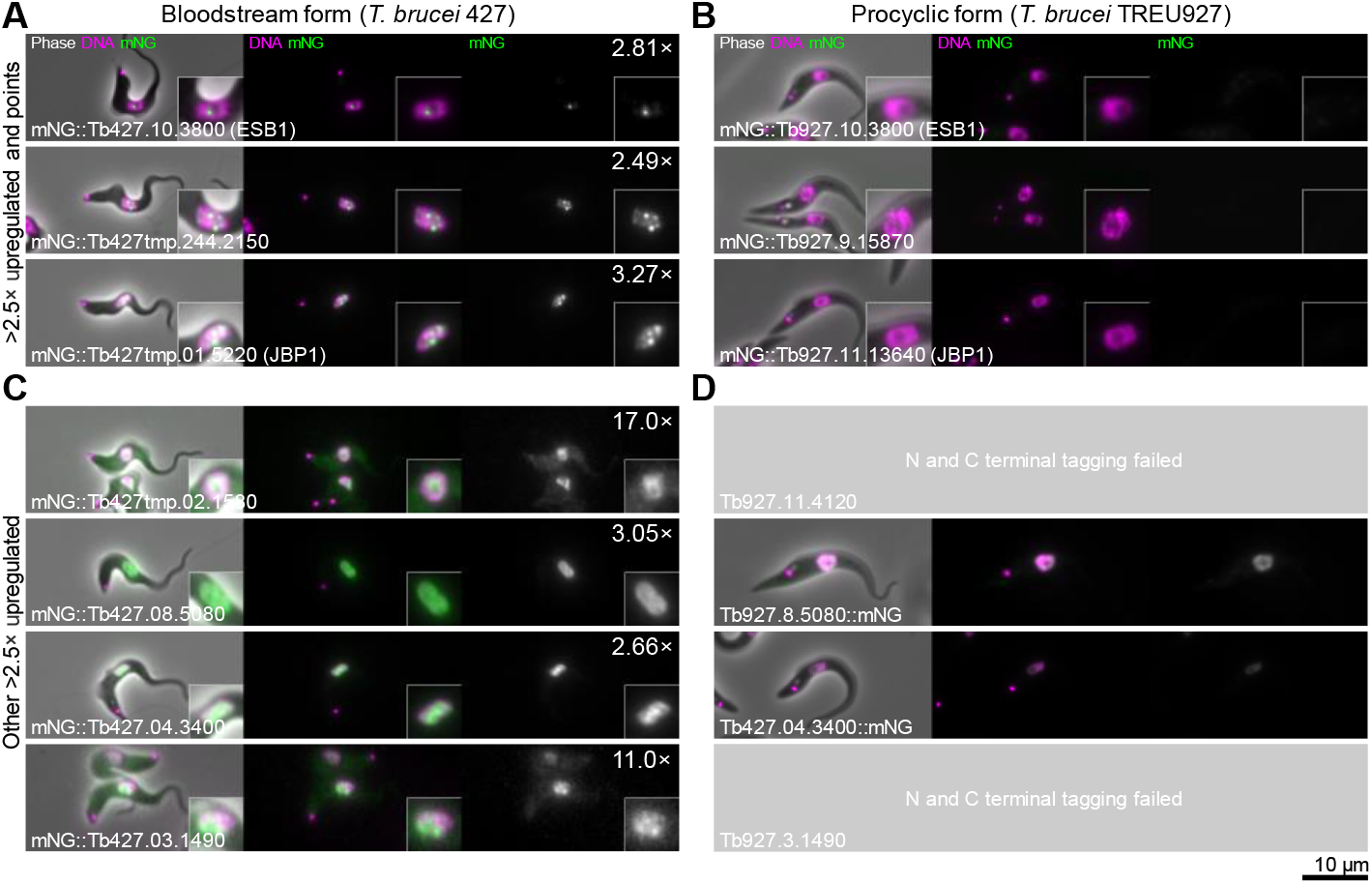
Bloodstream form-upregulated nuclear *T. brucei* proteins. Epifluorescence images of cell lines expressing tagged proteins. Images for each cell line are laid in the same format: Left, an overlay of the phase contract (grey), mNG fluorescence (green) and Hoechst DNA stain (magenta). Middle, the DNA stain and the mNG fluorescence. Right, the mNG fluorescence in greyscale. Fold upregulation in bloodstream form cells is shown in the top right. **A)** Subcellular localisation of all 3 proteins >2.5× upregulated in bloodstream forms^43^ which localised to one or multiple points in the nucleus when tagged at the N terminus in bloodstream forms. **B)** Subcellular localisation of the 3 proteins in A) in equivalent procyclic form cell lines, shown at the same contrast levels. **C)** Subcellular localisation of the remaining 4 proteins >2.5× upregulated in bloodstream forms which localised to the wider nucleus when tagged at the N terminus in bloodstream forms. **D)** Subcellular localisation of the 4 proteins in C) in equivalent procyclic form cell lines, shown at the same contrast levels. We were unable to generate two cell lines.

**Extended data fig. 2.**
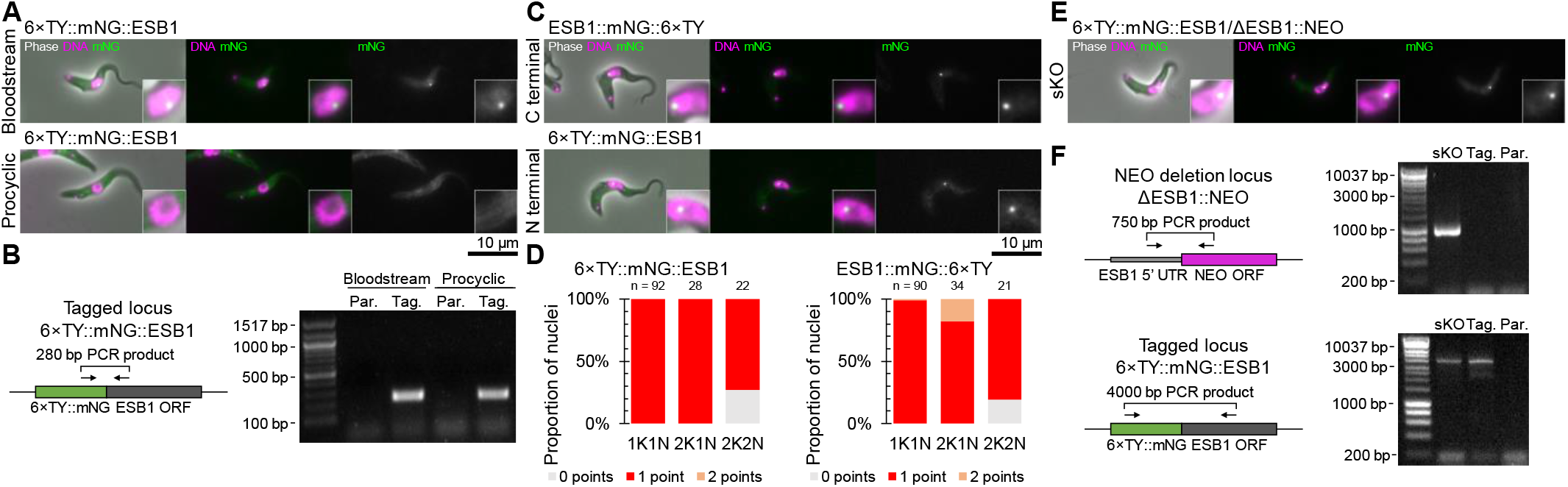
Tagging does not perturb ESB1 localisation or function. **A)** Clonal bloodstream form and procyclic form cell lines expressing Tb427.10.3800 or Tb927.10.3800 (ESB1) N terminally tagged with 6×TY::mNG respectively were re-generated following the initial screen. Epifluorescence images of the localisation of the tagged protein by mNG fluorescence. **B)** Confirmation of the expected genetic modification of the cell lines in A) by PCR from genomic DNA using a forward mNG and a reverse ESB1 ORF primer. Schematic shows the primer binding sites, DNA gel shows the resulting PCR products from extracted genomic DNA from the tagged (Tag.) or parental (Par.) cell line. **C)** Epifluorescence images of bloodstream form cell lines expressing 6×TY::mNG::ESB1 (N terminal tag), ESB1::mNG::6×TY (C terminal tag). **D)** Count of the number of points per nucleus at different stages of the cell cycle (1K1N, 2K1N and 2K2N) for N or C terminally tagged ESB1. **E)** Epifluorescence image of a single knockout (sKO) bloodstream form cell line with one N terminally tagged ESB1 allele and the other deleted by replacement with a drug selectable marker. **F)** PCR validation of the sKO cell line. Schematics represent the deleted ESB1 ORF (top) and N terminally tagged locus (bottom) and primer binding sites, DNA gels shows the resulting PCR products from extracted genomic DNA.

**Extended data fig. 3.**
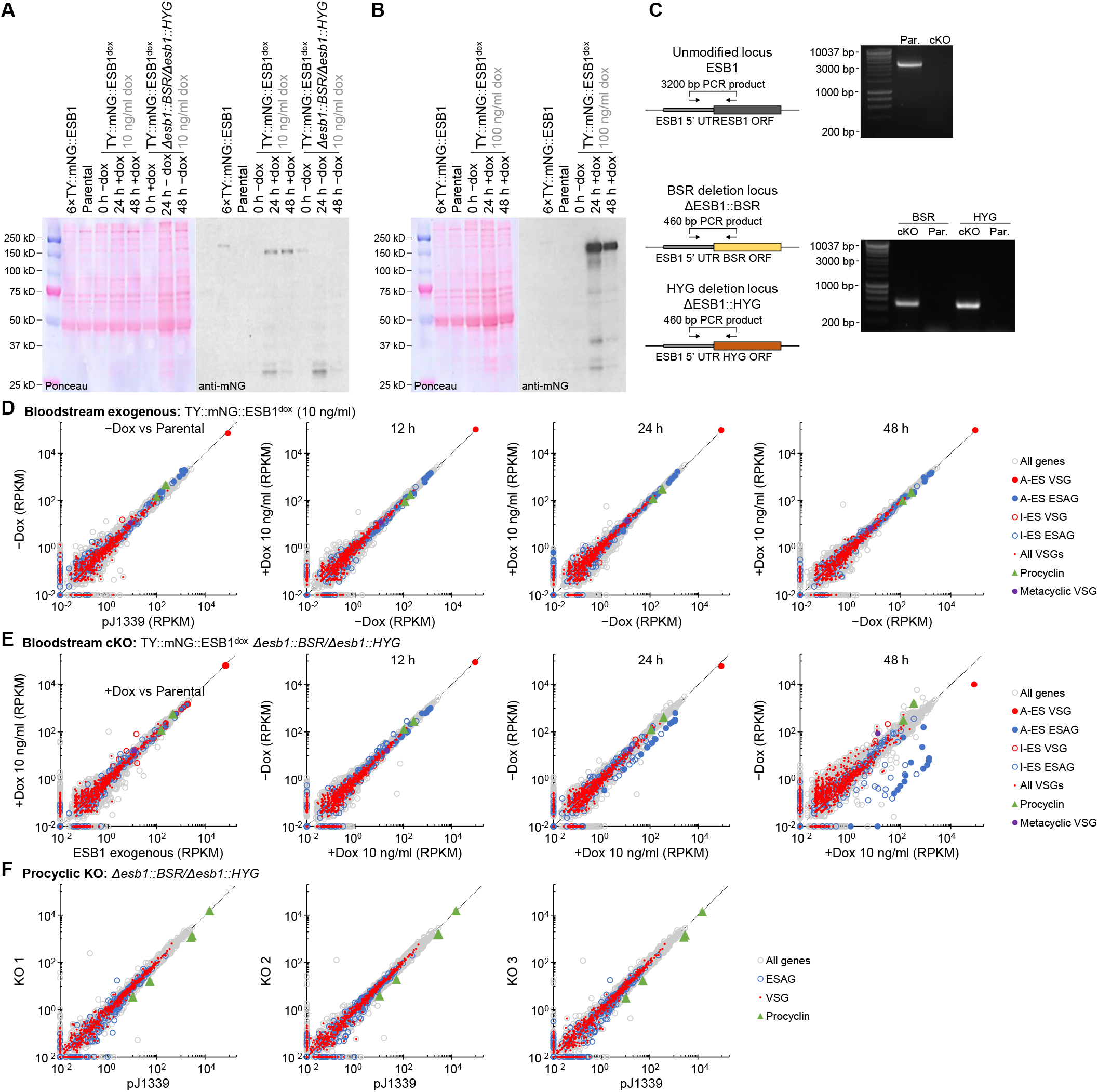
Generation and validation of an ESB1 conditional knockout. **A)** Western blot validation of the cKO cell line and the BSF pDex577 tagged ESB1 exogenous expression cell line (the intermediate in cKO generation), both induced with 10 ng/ml doxycycline. Predicted molecular weights for ESB1 are: 108 kDa (untagged), 137 kDa (TY::mNG tag) and 145 kDa (6×TY::mNG tag). The ponceau-stained membrane and anti-mNG blot are shown. **B)** Western blot validation of the BSF pDex577 tagged ESB1 exogenous expression cell line induced with 100 ng/ml doxycycline for overexpression. **C)** mRNA abundance in the BSF pDex577 tagged ESB1 exogenous expression cell line, plotting RPKM of uniquely mapped RNAseq reads. The exogenous expression prior to addition of doxycycline (0 h) is plotted relative to the parental pJ1339 cell line. Other plots are 12, 24 and 48 h after addition of 10 ng/ml doxycycline relative to the cell line grown without doxycycline. **D)** Validation of genetic modifications of the ESB1 conditional knockout (cKO). Schematics represent the deleted and tagged loci and primer binding sites and orientations, DNA gels shows the resulting PCR products from extracted genomic DNA. **E)** mRNA abundance in the cKO, plotting RPKM of uniquely mapped RNAseq reads. The cKO prior to doxycycline washout (0 h) is plotted relative to the BSF pDex577 tagged ESB1 exogenous expression cell line induced with 100 ng/ul doxycycline. Other plots are 12, 24 and 48 h after doxycycline washout relative to the cell line grown with 100 ng/ml doxycycline. **F)** mRNA abundance in the procyclic form KO, plotting RPKM of uniquely mapped RNAseq reads for three clonal KO cell lines relative to the parental cell line.

**Extended data fig. 4.**
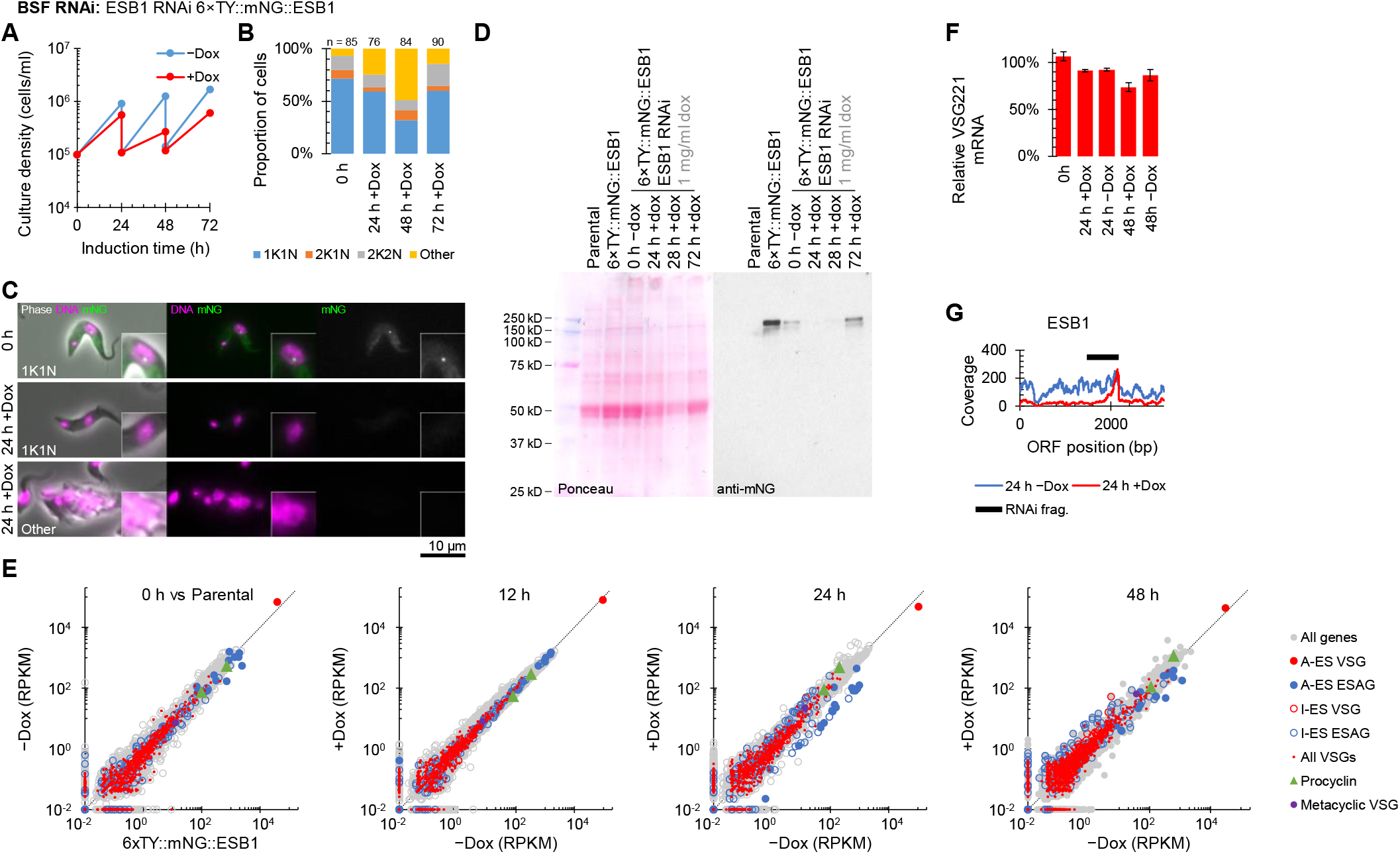
RNAi knockdown confirms ESB1 is vital and required for active BES transcription. Cellular phenotype of ESB1 RNAi knockdown. **A)** Growth curve of the ESB1 RNAi following doxycycline induction (+Dox) in comparison to uninduced (−Dox), using repeated subculture to maintain culture density under ∼1×10^6^ cells/ml. **B)** Number of kinetoplasts (K) and nuclei (N) per cell, counted by light microscopy, at 24 h intervals following washout of doxycycline from the ESB1 cKO. 1K1N, 2K1N and 2K2N are normal cell cycle stages. **C)** Representative epifluorescence microscope images of the ESB1 RNAi cell line before and after induction, showing a morphologically normal (1K1N) cell and a typical abnormal cell after 24 h induction. mNG signal is not detectable after 24 h induction. **D)** Anti-mNG Western blot validation of the ESB1 RNAi cell line. **E)** mRNA abundance in the BSF ESB1 RNAi cell line, plotting RPKM of uniquely mapped RNAseq reads. The uninduced cell line (0 h) is plotted relative to the parental 6×TY::mNG::ESB1 cell line. Other plots are 12, 24 and 48 h after addition of 1 mg/ml doxycycline relative to the cell line grown without doxycycline. **F)** qRT-PCR measurement of active BES VSG mRNA relative to the parental cell line. **G)** RNAseq read coverage over the ESB1 open reading frame shows reduced ESB1 transcript.

**Extended data fig. 5.**
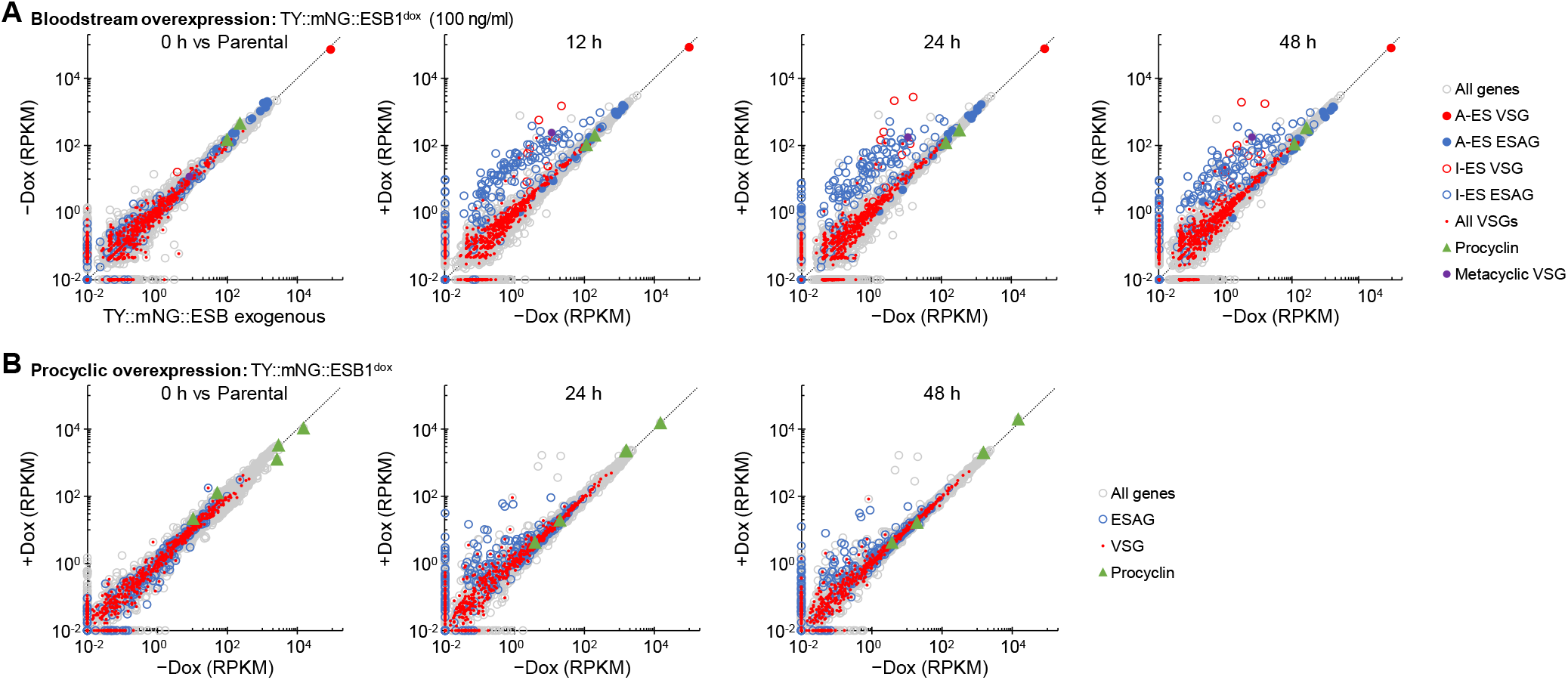
Extended analysis of the bloodstream and procyclic form overexpression analysis. **A)** mRNA abundance in the bloodstream form ESB1 overexpression cell line, plotting RPKM of uniquely mapped RNAseq reads. The overexpression prior to doxycycline addition (0 h) is plotted relative to the parental pJ1339 cell line. Other plots are 12, 24 and 48 h after 100 ng/ml doxycycline addition relative to the cell line grown without doxycycline. **B)** mRNA abundance in the procyclic form ESB1 overexpression cell line, plotting RPKM of uniquely mapped RNAseq reads. The overexpression prior to doxycycline addition (0 h) is plotted relative to the parental pJ1339 cell line. Other plots are 12, 24 and 48 h after 1 mg/ml doxycycline addition relative to the cell line grown without doxycycline.

## References

1. Vickerman, K. On the surface coat and flagellar adhesion in trypanosomes. Journal of cell science 5, 163–193 (1969).

2. Pays, E. et al. The genes and transcripts of an antigen gene expression site from T. brucei. Cell 57, 835–845 (1989).

3. Navarro, M. & Gull, K. A pol I transcriptional body associated with VSG mono-allelic expression in Trypanosoma brucei. Nature 414, 759–763 (2001).

4. Cully, D. F., Ip, H. S. & Cross, G. A. M. Coordinate transcription of variant surface glycoprotein genes and an expression site associated gene family in Trypanosoma brucei. Cell 42, 173–182 (1985).

5. Hertz-Fowler, C. et al.. Telomeric Expression Sites Are Highly Conserved in Trypanosoma brucei. PLOS ONE 3, e3527 (2008).

6. Xong, H. V. et al.. A VSG Expression Site–Associated Gene Confers Resistance to Human Serum in Trypanosoma rhodesiense. Cell 95, 839–846 (1998).

7. Cross, G. A. M., Kim, H.-S. & Wickstead, B. Capturing the variant surface glycoprotein repertoire (the VSGnome) of Trypanosoma brucei Lister 427. Mol. Biochem. Parasitol. 195, 59–73 (2014).

8. Müller, L. S. M. et al.. Genome organization and DNA accessibility control antigenic variation in trypanosomes. Nature 563, 121 (2018).

9. Benmerzouga, I. et al.. Trypanosoma brucei Orc1 is essential for nuclear DNA replication and affects both VSG silencing and VSG switching. Molecular Microbiology 87, 196–210 (2013).

10. Jehi, S. E., Nanavaty, V. & Li, B. Trypanosoma brucei TIF2 and TRF Suppress VSG Switching Using Overlapping and Independent Mechanisms. PLOS ONE 11, e0156746 (2016).

11. Reynolds, D. et al.. Regulation of transcription termination by glucosylated hydroxymethyluracil, base J, in Leishmania major and Trypanosoma brucei. Nucleic Acids Res 42, 9717–9729 (2014).

12. Yang, X., Figueiredo, L. M., Espinal, A., Okubo, E. & Li, B. RAP1 is essential for silencing telomeric variant surface glycoprotein genes in Trypanosoma brucei. Cell 137, 99–109 (2009).

13. Alsford, S. & Horn, D. Cell-cycle-regulated control of VSG expression site silencing by histones and histone chaperones ASF1A and CAF-1b in Trypanosoma brucei. Nucleic Acids Res 40, 10150–10160 (2012).

14. Denninger, V. & Rudenko, G. FACT plays a major role in histone dynamics affecting VSG expression site control in Trypanosoma brucei. Mol Microbiol 94, 945–962 (2014).

15. Figueiredo, L. M., Janzen, C. J. & Cross, G. A. M. A Histone Methyltransferase Modulates Antigenic Variation in African Trypanosomes. PLOS Biology 6, e161 (2008).

16. Hughes, K. et al.. A novel ISWI is involved in VSG expression site downregulation in African trypanosomes. EMBO J 26, 2400–2410 (2007).

17. Narayanan, M. S. et al.. NLP is a novel transcription regulator involved in VSG expression site control in Trypanosoma brucei. Nucleic Acids Res 39, 2018–2031 (2011).

18. Schulz, D. et al.. Bromodomain Proteins Contribute to Maintenance of Bloodstream Form Stage Identity in the African Trypanosome. PLOS Biology 13, e1002316 (2015).

19. Wang, Q.-P., Kawahara, T. & Horn, D. Histone deacetylases play distinct roles in telomeric VSG expression site silencing in African trypanosomes. Mol Microbiol 77, 1237–1245 (2010).

20. Narayanan, M. S. & Rudenko, G. TDP1 is an HMG chromatin protein facilitating RNA polymerase I transcription in African trypanosomes. Nucleic Acids Res 41, 2981–2992 (2013).

21. López-Farfán, D., Bart, J.-M., Rojas-Barros, D. I. & Navarro, M. SUMOylation by the E3 ligase TbSIZ1/PIAS1 positively regulates VSG expression in Trypanosoma brucei. PLoS Pathog. 10, e1004545 (2014).

22. Saura, A. et al.. SUMOylated SNF2PH promotes variant surface glycoprotein expression in bloodstream trypanosomes. EMBO Rep. 20, e48029 (2019).

23. Glover, L., Hutchinson, S., Alsford, S. & Horn, D. VEX1 controls the allelic exclusion required for antigenic variation in trypanosomes. Proc. Natl. Acad. Sci. U.S.A. 113, 7225–7230 (2016).

24. Faria, J. et al.. Monoallelic expression and epigenetic inheritance sustained by a Trypanosoma brucei variant surface glycoprotein exclusion complex. Nature Communications 10, 1–14 (2019).

25. Faria, J. et al.. Spatial integration of transcription and splicing in a dedicated compartment sustains monogenic antigen expression in African trypanosomes. Nature Microbiology 1–12 (2021) doi:10.1038/s41564-020-00833-4.

26. Boothroyd, J. C. & Cross, G. A. M. Transcripts coding for variant surface glycoproteins of Trypanosoma brucei have a short, identical exon at their 5′ end. Gene 20, 281–289 (1982).

27. Daniels, J.-P., Gull, K. & Wickstead, B. Cell biology of the trypanosome genome. Microbiol. Mol. Biol. Rev. 74, 552–569 (2010).

28. Dean, S., Sunter, J. D. & Wheeler, R. J. TrypTag.org: A Trypanosome Genome-wide Protein Localisation Resource. Trends in Parasitology 33, 80–82 (2017).

29. Landeira, D., Bart, J.-M., Van Tyne, D. & Navarro, M. Cohesin regulates VSG monoallelic expression in trypanosomes. J. Cell Biol. 186, 243–254 (2009).

30. Fadda, A. et al.. Transcriptome-wide analysis of trypanosome mRNA decay reveals complex degradation kinetics and suggests a role for co-transcriptional degradation in determining mRNA levels. Mol Microbiol 94, 307–326 (2014).

31. Ullu, E., Tschudi, C. & Chakraborty, T. RNA interference in protozoan parasites. Cell. Microbiol. 6, 509–519 (2004).

32. LaCount, D. J. et al.. Analysis of a donor gene region for a variant surface glycoprotein and its expression site in African trypanosomes. Nucleic Acids Res 29, 2012–2019 (2001).

33. Jayaraman, S. et al.. Application of long read sequencing to determine expressed antigen diversity in Trypanosoma brucei infections. PLOS Neglected Tropical Diseases 13, e0007262 (2019).

34. Budzak, J. et al.. Dynamic colocalization of 2 simultaneously active VSG expression sites within a single expression-site body in Trypanosoma brucei. Proc. Natl. Acad. Sci. U.S.A. 116, 16561–16570 (2019).

35. Nguyen, T. N., Müller, L. S. M., Park, S. H., Siegel, T. N. & Günzl, A. Promoter occupancy of the basal class I transcription factor A differs strongly between active and silent VSG expression sites in Trypanosoma brucei. Nucleic Acids Res. 42, 3164–3176 (2014).

## Additional references

36. Becker, M. et al.. Isolation of the repertoire of VSG expression site containing telomeres of Trypanosoma brucei 427 using transformation-associated recombination in yeast. Genome Res 14, 2319–2329 (2004).

37. Hirumi, H. & Hirumi, K. Continuous cultivation of Trypanosoma brucei blood stream forms in a medium containing a low concentration of serum protein without feeder cell layers. J. Parasitol. 75, 985–989 (1989).

38. Berriman, M. et al.. The genome of the African trypanosome Trypanosoma brucei. Science 309, 416–22 (2005).

39. Halliday, C. et al.. Cellular landmarks of Trypanosoma brucei and Leishmania mexicana. Molecular and Biochemical Parasitology (2018) doi:10.1016/j.molbiopara.2018.12.003.

40. Brun, R. & Schönenberger, M. Cultivation and in vitro cloning or procyclic culture forms of Trypanosoma brucei in a semi-defined medium. Short communication. Acta Trop 36, 289–292 (1979).

41. Alves, A. A. et al.. Control of assembly of extra-axonemal structures: the paraflagellar rod of trypanosomes. J Cell Sci 133, (2020).

42. Schumann Burkard, G., Jutzi, P. & Roditi, I. Genome-wide RNAi screens in bloodstream form trypanosomes identify drug transporters. Mol Biochem Parasitol 175, 91–94 (2011).

43. Jensen, B. C. et al.. Extensive stage-regulation of translation revealed by ribosome profiling of Trypanosoma brucei. BMC Genomics 15, 911 (2014).

44. Jensen, B. C., Sivam, D., Kifer, C. T., Myler, P. J. & Parsons, M. Widespread variation in transcript abundance within and across developmental stages of Trypanosoma brucei. BMC Genomics 10, 482 (2009).

45. Naguleswaran, A., Doiron, N. & Roditi, I. RNA-Seq analysis validates the use of culture-derived Trypanosoma brucei and provides new markers for mammalian and insect life-cycle stages. BMC Genomics 19, 227 (2018).

46. Siegel, T. N., Hekstra, D. R., Wang, X., Dewell, S. & Cross, G. A. M. Genome-wide analysis of mRNA abundance in two life-cycle stages of Trypanosoma brucei and identification of splicing and polyadenylation sites. Nucleic Acids Res 38, 4946–4957 (2010).

47. Vasquez, J.-J., Hon, C.-C., Vanselow, J. T., Schlosser, A. & Siegel, T. N. Comparative ribosome profiling reveals extensive translational complexity in different Trypanosoma brucei life cycle stages. Nucleic Acids Res 42, 3623–3637 (2014).

48. Beneke, T. et al.. A CRISPR Cas9 high-throughput genome editing toolkit for kinetoplastids. Open Science 4, 170095 (2017).

49. Dean, S. et al.. A toolkit enabling efficient, scalable and reproducible gene tagging in trypanosomatids. Open Biol 5, 140197 (2015).

50. Shaner, N. C. et al.. A bright monomeric green fluorescent protein derived from Branchiostoma lanceolatum. Nat Meth 10, 407–409 (2013).

51. Kelly, S. et al.. Functional genomics in Trypanosoma brucei: a collection of vectors for the expression of tagged proteins from endogenous and ectopic gene loci. Mol. Biochem. Parasitol. 154, 103–109 (2007).

52. Poon, S. K., Peacock, L., Gibson, W., Gull, K. & Kelly, S. A modular and optimized single marker system for generating Trypanosoma brucei cell lines expressing T7 RNA polymerase and the tetracycline repressor. Open Biology 2, 110037 (2012).

53. Bastin, P., Bagherzadeh, A., Matthews, K. R. & Gull, K. A novel epitope tag system to study protein targeting and organelle biogenesis in Trypanosoma brucei. Molecular and Biochemical Parasitology 77, 235–239 (1996).

54. Dean, S. & Sunter, J. Light Microscopy in Trypanosomes: Use of Fluorescent Proteins and Tags. Methods Mol. Biol. 2116, 367–383 (2020).

55. Edelstein, A., Amodaj, N., Hoover, K., Vale, R. & Stuurman, N. Computer control of microscopes using µManager. Curr Protoc Mol Biol Chapter 14, Unit14.20 (2010).

56. Wheeler, R. J. ImageJ for Partially and Fully Automated Analysis of Trypanosome Micrographs. in Trypanosomatids: Methods and Protocols (eds. Michels, P. A. M., Ginger, M. L. & Zilberstein, D.) 385–408 (Springer US, 2020). doi:10.1007/978-1-0716-0294-2_24.

57. Wheeler, R. J., Gull, K. & Gluenz, E. Detailed interrogation of trypanosome cell biology via differential organelle staining and automated image analysis. BMC Biology 10, 1 (2012).

58. Aslett, M. et al.. TriTrypDB: a functional genomic resource for the Trypanosomatidae. Nucleic Acids Res 38, D457–D462 (2010).

59. Warrenfeltz, S. et al.. EuPathDB: The Eukaryotic Pathogen Genomics Database Resource. Methods Mol Biol 1757, 69–113 (2018).

60. Weeks, A. F., Nathan. Best Practices for De Novo Transcriptome Assembly with Trinity. Harvard FAS Informatics ./best-practices-for-de-novo-transcriptome-assembly-with-trinity.html (2020).

61. Song, L. & Florea, L. Rcorrector: efficient and accurate error correction for Illumina RNA-seq reads. GigaScience 4, 48 (2015).

62. Nguyen, T. N., Nguyen, B. N., Lee, J. H., Panigrahi, A. K. & Günzl, A. Characterization of a Novel Class I Transcription Factor A (CITFA) Subunit That Is Indispensable for Transcription by the Multifunctional RNA Polymerase I of Trypanosoma brucei. Eukaryot Cell 11, 1573–1581 (2012).

63. Jehi, S. E., Wu, F. & Li, B. Trypanosoma brucei TIF2 suppresses VSG switching by maintaining subtelomere integrity. Cell Res 24, 870–885 (2014).

64. Jehi, S. E. et al.. Suppression of subtelomeric VSG switching by Trypanosoma brucei TRF requires its TTAGGG repeat-binding activity. Nucleic Acids Res 42, 12899–12911 (2014).

65. Davies, C. et al.. TbSAP is a novel chromatin protein repressing metacyclic variant surface glycoprotein expression sites in bloodstream form Trypanosoma brucei. Nucleic Acids Res (2021) doi:10.1093/nar/gkab109.

66. DuBois, K. N. et al.. NUP-1 Is a Large Coiled-Coil Nucleoskeletal Protein in Trypanosomes with Lamin-Like Functions. PLOS Biology 10, e1001287 (2012).

67. Maishman, L. et al.. Co-dependence between trypanosome nuclear lamina components in nuclear stability and control of gene expression. Nucleic Acids Res 44, 10554–10570 (2016).

68. Aresta-Branco, F., Sanches-Vaz, M., Bento, F., Rodrigues, J. A. & Figueiredo, L. M. African trypanosomes expressing multiple VSGs are rapidly eliminated by the host immune system. Proc Natl Acad Sci U S A 116, 20725–20735 (2019).

69. Povelones, M. L., Gluenz, E., Dembek, M., Gull, K. & Rudenko, G. Histone H1 Plays a Role in Heterochromatin Formation and VSG Expression Site Silencing in Trypanosoma brucei. PLOS Pathogens 8, e1003010 (2012).

70. Black, J. A. et al.. Trypanosoma brucei ATR Links DNA Damage Signaling during Antigenic Variation with Regulation of RNA Polymerase I-Transcribed Surface Antigens. Cell Reports 30, 836-851.e5 (2020).

71. Cestari, I. & Stuart, K. Inositol phosphate pathway controls transcription of telomeric expression sites in trypanosomes. PNAS 112, E2803–E2812 (2015).

72. Cestari, I., McLeland-Wieser, H. & Stuart, K. Nuclear Phosphatidylinositol 5-Phosphatase Is Essential for Allelic Exclusion of Variant Surface Glycoprotein Genes in Trypanosomes. Molecular and Cellular Biology 39, (2019).

